# Rare variant enriched identity-by-descent enables the detection of distant relatedness and older divergence between populations

**DOI:** 10.1101/2020.05.05.079541

**Authors:** Amol C. Shetty, NHLBI Trans-Omics for Precision Medicine (TOPMed) Consortium, TOPMed Population Genetics Working Group, Jeffrey O’Connell, Braxton D. Mitchell, Timothy D. O’Connor

## Abstract

**Motivation:** The global human population has experienced an explosive growth from a few million to roughly 7 billion people in the last 10,000 years. Accompanying this growth has been the accumulation of rare variants that can inform our understanding of human evolutionary history. Common variants have primarily been used to infer the structure of the human population and relatedness between two individuals. However, with the increasing abundance of rare variants observed in large-scale projects, such as Trans-Omics for Precision Medicine (TOPMed), the use of rare variants to decipher cryptic relatedness and fine-scale population structure can be beneficial to the study of population demographics and association studies. Identity-by-descent (IBD) is an important framework used for identifying these relationships. IBD segments are broken down by recombination over time, such that longer shared haplotypes give strong evidence of recent relatedness while shorter shared haplotypes are indicative of more distant relationships. Current methods to identify IBD accurately detect only long segments (> 2cM) found in related individuals.

**Algorithm:** We describe a metric that leverages rare-variants shared between individuals to improve the detection of short IBD segments. We computed IBD segments using existing methods implemented in *Refined IBD* where we enrich the signal using our metric that facilitates the detection of short IBD segments (<2cM) by explicitly incorporating rare variants.

**Results:** To test our new metric, we simulated datasets involving populations with varying divergent time-scales. We show that rare-variant IBD identifies shorter segments with greater confidence and enables the detection of older divergence between populations. As an example, we applied our metric to the Old-Order Amish cohort with known genealogies dating 14 generations back to validate its ability to detect genetic relatedness between distant relatives. This analysis shows that our method increases the accuracy of identifying shorter segments that in turn capture distant relationships.

**Conclusions:** We describe a method to enrich the detection of short IBD segments using rare-variant sharing within IBD segments. Leveraging rare-variant sharing improves the information content of short IBD segments better than common variants alone. We validated the method in both simulated and empirical datasets. This method can benefit association analyses, IBD mapping analyses, and demographic inferences.

## Introduction

Relatedness is the estimation of shared ancestry between individuals and is a fundamental concept in genetics that plays an integral role in many fields ranging from animal breeding, human disease gene mapping, and forensic science. For example, studies involving population structure (Pritchard et al. 2000; Novembre et al. 2008; Biswas et al. 2009; Lawson et al. 2012; Elhaik et al. 2014) make inferences about shared ancestry based on estimates of genetic relatedness. Population structure often closely agrees with geography under the assumption that populations can be partitioned into “islands” of individuals with increased intra-island mating and decreased inter-island migrations (Astle and Balding 2009). Seminal work by Novembre and colleagues (Novembre et al. 2008) showed that genetic markers mirror geography in Europe allowing for accurate inference of geographic origin using an individual’s DNA.

Failure to estimate cryptic relatedness among study subjects in genetic association studies can inflate statistical evidence for association. Relatedness estimates are thus integral to the performance of genetic association studies that have traditionally been estimated from pedigrees and more recently from genomic data in large populations lacking pedigree information (Purcell et al. 2007; Manichaikul et al. 2010; Thornton et al. 2012). Mixed linear models are commonly used in genetic association studies in order to incorporate a genetic relationship matrix (GRM) that help to prevent false-positive associations due to population structure or relatedness (Kang et al. 2010; Price et al. 2010; Zhang et al. 2010). More recently, commercial products such as those provided by AncestryDNA™, use genetic information to identify recent relatedness between its consumers allowing them to discern distant relatives who may not be represented in their documented pedigrees (Han et al. 2017). Similarly, the field of forensic science has utilized large public repositories to identify relatives of forensic samples since the alleged suspect may share genetic information inherited from a common ancestor with relatives represented in these repositories (Jobling and Gill 2004; Weir et al. 2006; Kayser and de Knijff 2011).

Genetic relatedness between two individuals can be defined as the probability that their alleles were inherited from a common ancestor, in other words the alleles are identical-by-descent (IBD). Traditionally, relatedness was estimated directly from known pedigrees and was limited by the generational depth of the pedigree. However, datasets from modern sampling procedures are often not accompanied with any pedigree information or have incomplete pedigrees (Wellcome Trust Case Control Consortium 2007; International HapMap Consortium et al. 2007; Auton et al. 2009; Henn et al. 2010; Ralph and Coop 2013; Carlson et al. 2018). With the introduction of high-throughput genotyping and sequencing technologies, the relatedness estimates can be made directly from genotype information without prior information about the genealogy of the samples.

Over the past decade, multiple methods have been developed to estimate relatedness from single nucleotide polymorphism (SNP) information. Software tools such *PLINK* (Purcell et al. 2007; Chang et al. 2015), *KING* (Manichaikul et al. 2010), *REAP* (Thornton et al. 2012), and *PC-Relate* (Conomos et al. 2016) use allele frequencies across multiple loci to estimate the probabilities of sharing zero, one, or two alleles at a given genetic locus corresponding to the number of copies that are in IBD. When summed across multiple loci these estimates can be transformed into a kinship coefficient estimate for a pair of individuals. Allele frequency-based kinship estimates utilize genome-wide averages of single-SNP statistics that do not take the lengths of genomic regions shared between two individuals into account. However, software tools like *Germline* (Gusev et al. 2009), *Beagle* (Browning and Browning 2010), *fastIBD* (Browning and Browning 2011), and *Refined IBD* (Browning and Browning 2013b) detect IBD segments in the genome shared between pairs of individuals that have been used to infer the degree of relatedness between individuals. More recently, the *KING* software has incorporated IBD-based kinship estimates.

Alternatively, other software tools such as *RelateAdmix* (Moltke and Albrechtsen 2014), *ERSA* (Huff et al. 2011; Li et al. 2014), and *HaploScore* (Durand et al. 2014) combine multiple statistics to improve these estimates of relatedness. *RelateAdmix* uses a maximum likelihood estimator that combines allele frequencies with admixture proportions while *ERSA* utilizes a maximum likelihood method to estimate recent shared ancestry from the number and length of IBD segments. *HaploScore* takes a different approach in order to improve the accuracy of segments detected as IBD by incorporating different error statistics. All of the above methods have high accuracy when detecting first-degree to third-degree relationships (Ramstetter et al. 2017) and haplotype-based methods perform better than allele frequency-based methods (Gattepaille and Jakobsson 2012; Gusev et al. 2012; Palamara et al. 2012). However, other software tools such as *PRIMUS* (Staples et al. 2014) and *PADRE* (Staples et al. 2016) were developed to estimate more distant relationships by reconstructing pedigrees from existing relatedness estimates.

The scale of human sequencing studies has grown exponentially in recent years with the initiation of large-scale projects such as DiscovEHR (Dewey et al. 2016), UK Biobank (Sudlow et al. 2015), Precision Medicine Initiative (Collins and Varmus 2015), TOPMed (Taliun et al. 2019), MVP (Gaziano et al. 2016), and the All of Us Research Program that involve hundreds of thousands of samples. These studies include large number of related individuals between whom cryptic relatedness can be uncovered using their genetic data. For example, nearly 30% of the UK Biobank participants were found to be related (3^rd^ order or closer) to another person in the cohort (Bycroft et al. 2018). Estimates of cryptic relatedness would help in building a better GRM and in turn improve analyses that are sensitive to inaccurate estimates of relationships and incorrect pedigrees in large-scale genetic association cohorts. A feature of large-scale human sequencing projects is the presence of numerous rare variants with minor allele frequencies as low as 1 x 10^−5^. Rare variants have typically arisen as de-novo mutations concurrent with the explosion of population size in recent generations (Keinan and Clark 2012). Their recent origins compared to common variants make them a powerful resource for delineating fine-scale population structure (Baye et al. 2011; Keinan and Clark 2012; Tennessen et al. 2012; O’Connor et al. 2015).

IBD segments resulting from shared ancestry are detected mainly as large segments (2cM and larger) with greater accuracy of detection as the length increases (Chapman and Thompson 2003; Moltke et al. 2011; Gusev et al. 2009; Browning and Browning 2013b; Ralph and Coop 2013). Current haplotype sharing methods for estimation of relatedness use common variants (minor allele frequency > 5%) since phasing of variants into haplotypes is critical to IBD detection and it is difficult to phase rare variants. Hence, these methods have the disadvantage of not using information available from the sharing of rare variants between two individuals. Since the underlying principle of IBD detection is based on the sharing of very low frequency haplotypes, rare variants can be highly informative for IBD detection. While common variants can be shared between two individuals by chance without the presence of a common ancestor, sharing of rare variants between two individuals without a common ancestor is less likely. Hence, shared rare variants provide additional evidence of haplotype sharing that will increase the odds of IBD vs identity-by-state (IBS) due to rarity of co-ancestry and in turn improve the accuracy of IBD detection especially for short segments with lengths less than 2cM.

In the *Algorithm and Implementation* section, we describe a novel method for enriching the detection of haplotype sharing between two individuals by leveraging shared rare variants. In the *Results* section, we evaluate the method in simulated datasets as well as an empirical dataset involving a 14-generation founder population of Old Order Amish (OOA) individuals, 1,100 of whom were sequenced as part of the TOPMed initiative (Mitchell et al. 2008; Taliun et al. 2019). We then describe the assessment of the rare-variant IBD metric (rvIBD) in the simulated dataset and evaluate its performance in detecting higher-order relatedness using prior knowledge about the different degrees of relationships ascertained from the OOA pedigree (Agarwala et al. 1998).

## Algorithm and Implementation

We present the rare-variant IBD (rvIBD) metric, which identifies short IBD segments by leveraging shared rare variants between individuals from a large sequencing cohort. The implementation of the rvIBD metric involves three stages (<u>Supplementary Figure S1</u>). The first stage is the detection of IBD segments using common variants and existing methods such as Refined IBD (Browning and Browning 2013b) with relaxed filtering criteria to allow for the detection of short IBD segments (<2cM). The second stage is the identification of rare variants with minor allele frequency < 1%, that are shared between two samples. The third stage utilizes the combination of rare variant genotype and allele frequency within a Bayesian framework to update the odds of distinguishing IBD from IBS.

### Rare-variant IBD (rvIBD) Metric

The first step involves the detection of IBD segments between all pairs of samples from large-scale cohorts that include participants who have been sequenced or densely genotyped. We used Refined IBD (Browning and Browning 2013b), an accurate and computationally efficient algorithm, for IBD segment detection. Refined IBD uses a two-step process that first identifies shared haplotypes greater than a user-defined length threshold and then evaluates the evidence for IBD using a hidden Markov model. Using markers with a minimum minor allele frequency of 5%, we identified IBD segments using default parameters except for lowering the length threshold and the log-odds threshold. The second step involves the identification of rare variants shared between two samples. Excluding singletons, bi-allelic markers are first filtered to retain variants with a maximum minor allele frequency of 1%. The lower bound of the allele count for rare variants could be increased to control for genotyping errors depending on the yield of the different filters. For each IBD segment identified between two samples, we extract the genotypes for the rare variants within the endpoints of the segment. Finally, in the third step, we iterate over the set of rare variants and estimate the rvIBD metric by enriching the original log-odds (LOD) score from Refined IBD using the following equation based on the Bayesian approach for odds ratios,

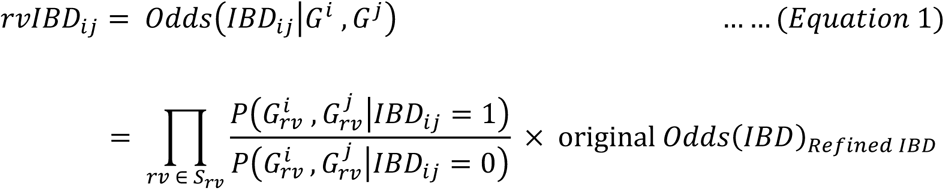

where,

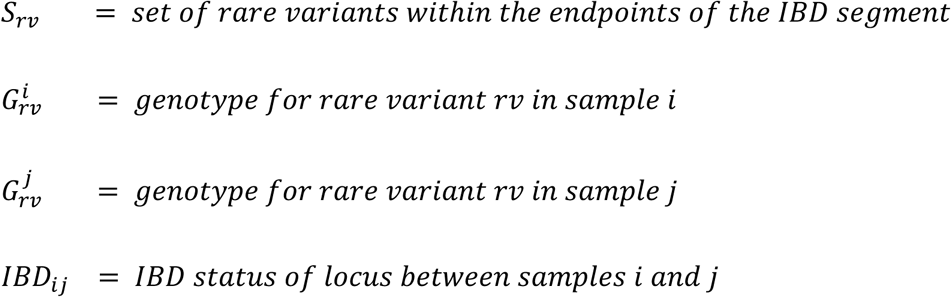

The probabilities for a pair of genotypes 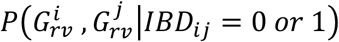 given IBD or non-IBD status are summarized in Figure 1. In order to incorporate allele error into the rvIBD metric, we replace the observed minor allele frequency *f*_*B*_ with the corrected minor allele frequency *p*_*B*_ = (*f*_*B*_ − *ε*)/(1 − 2*ε*) and corrected major allele frequency *p*_*A*_ = (1 − *p*_*B*_) where *ε* is the allele error at the biallellic marker with major allele A and minor allele B.

**Figure 1:**
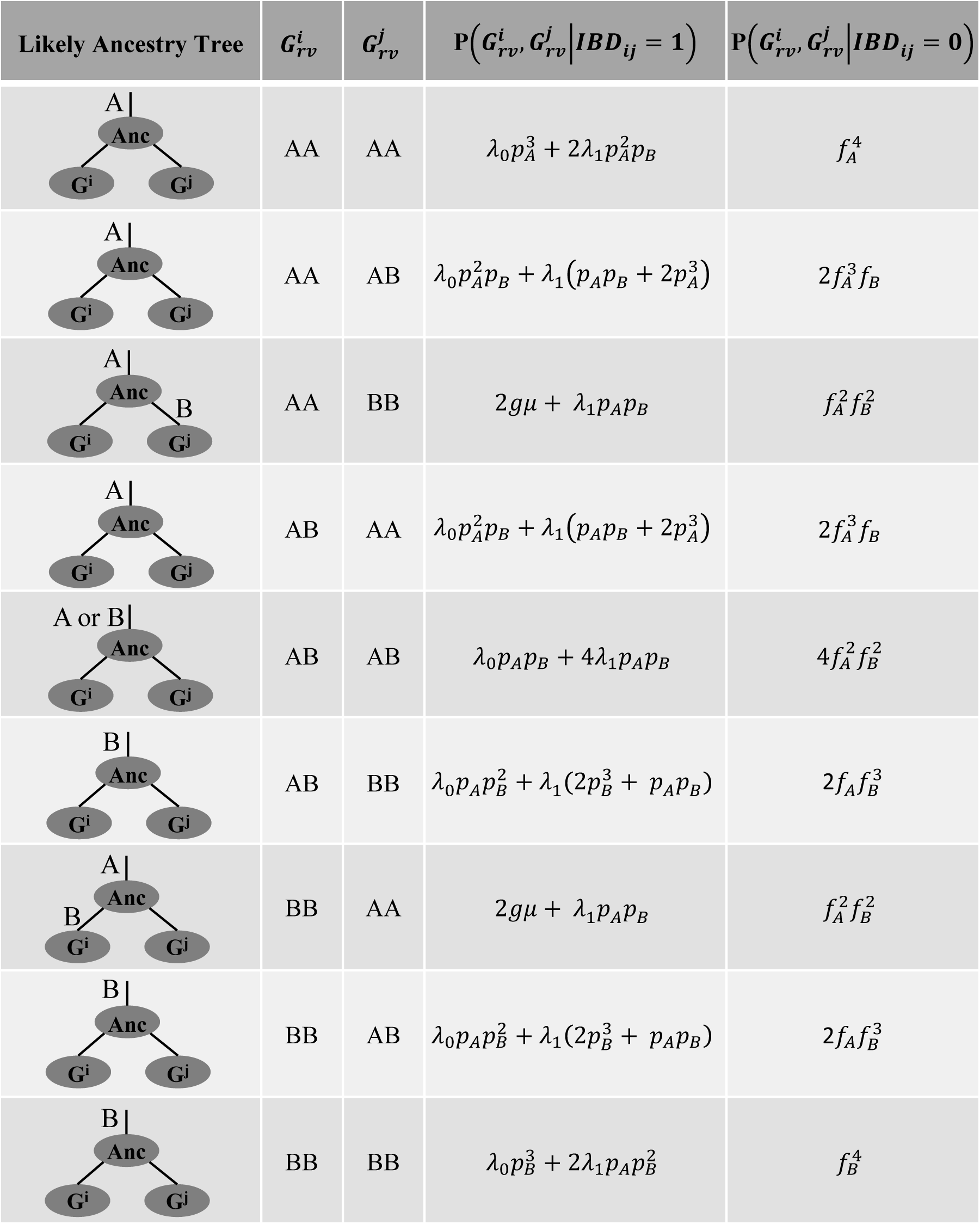
Probability of rare variant genotype pairs given IBD states. Probabilities for rare-variant genotype pairs with major allele A and minor allele B are computed using corrected allele frequencies given by *p*_*A*_ and *p*_*B*_ respectively and adjusted for allele error *ε* using error term *λ*_*j*_ = *ε*^*j*^(1 − *ε*)^(4−*j*)^ for *j* ∈ [0,1] (Browning and Browning 2013a). 2*gμ* is the probability of mutation since common ancestor *g* generations ago with a mutation rate *μ* (Palamara et al. 2015). *g* ≅ 3/2*l* where *l* is IBD segment length in Morgans (Baharian et al. 2016).

We also adjusted the probabilities for genotype pairs for both allele errors and mutations acquired since most recent common ancestor (Palamara et al. 2015; Baharian et al. 2016; Smith et al. 2018). Given that rare variants cannot be accurately phased, the rvIBD metric was designed specifically to use the rare variant genotype and the number of rare alleles shared between two individuals to enrich the odds of IBD vs non-IBD. The new rvIBD metric and number of shared rare variants can be used to assess accuracy of IBD detection, especially for short IBD segments.

## Results

In order to assess the performance of the rare-variant IBD metric (rvIBD), we evaluated its ability to detect short IBD segments between populations with older divergence using a simulated dataset. In addition, we applied the rvIBD metric to the Old Order Amish (OOA) cohort to assess detectability of distant relatedness using prior knowledge about the different degrees of relationships ascertained from the OOA pedigree (Agarwala et al. 1998).

### Simulated dataset

We simulated a demographic model using the program *msprime* (Kelleher et al. 2016), a coalescent-based simulator. Our estimated demographic model started with an initial population size of 12,300 about 220,000 ya, with an out-of-Africa reduction to 2,100 occurring 140,000 ya (Gutenkunst et al. 2009). This population then diverged into two populations of size 1000 and 510 respectively about T_POP2-POP3_ generations ago where T_POP2-POP3_ ranges from 5 generations (recent divergence) to 1000 generations ago (distant divergence) (<u>Supplementary Figure S2</u>). We used a mutation rate of 2 × 10^−8^ and a recombination rate of 2 × 10^−8^ (i.e. 1Mb ≅ 1cM). For each divergence time, we simulated 20 data sets consisting of 10 Mb segments and 1,500 diploid individuals (500 individuals per population). msprime outputs genotypes in VCF format as well as ancestry trees that can be queried to compute time to most recent common ancestor (TMRCA) for any genomic segment between any two individuals. Using markers with a minimum minor allele frequency of 5%, we identified IBD segments using default parameters except for lowering the length threshold to 0.15cM and the log-odds threshold to 1. Each IBD segment detected using *Refined IBD* was leveraged for the presence of shared rare variants (minor allele frequency < 1%) to compute the number of shared rare variants and the enriched rvIBD log-odds (rvLOD) score (see Equation 1). The IBD segments were further segregated in groups either based on the number of rare variants shared or based on the original LOD and enriched rvLOD scores. The distributions of the segment lengths were compared between groups using the Wilcoxon rank sum test (for 2 groups) and Kruskal-Wallis test (for 3 groups). We also estimated the TMRCA (in generations) for each IBD segment using the ancestry tree generated by msprime for each simulation (Kelleher et al. 2016). The distributions of the TMRCA estimates were also compared between groups based on the LOD and rvLOD scores using the Kruskal-Wallis test.

### Shared rare variants capable of tagging short IBD segments

For each simulation based on varying divergence times ranging from 5 to 1000 generations, we first binned the IBD segments into two categories, namely segments with no shared rare variants and segments with at least 1 shared rare variant within the endpoints of the segment. We then compared the distribution of IBD segment length between the two categories (Figure 2; <u>Supplementary Figure S3</u>).

**Figure 2:**
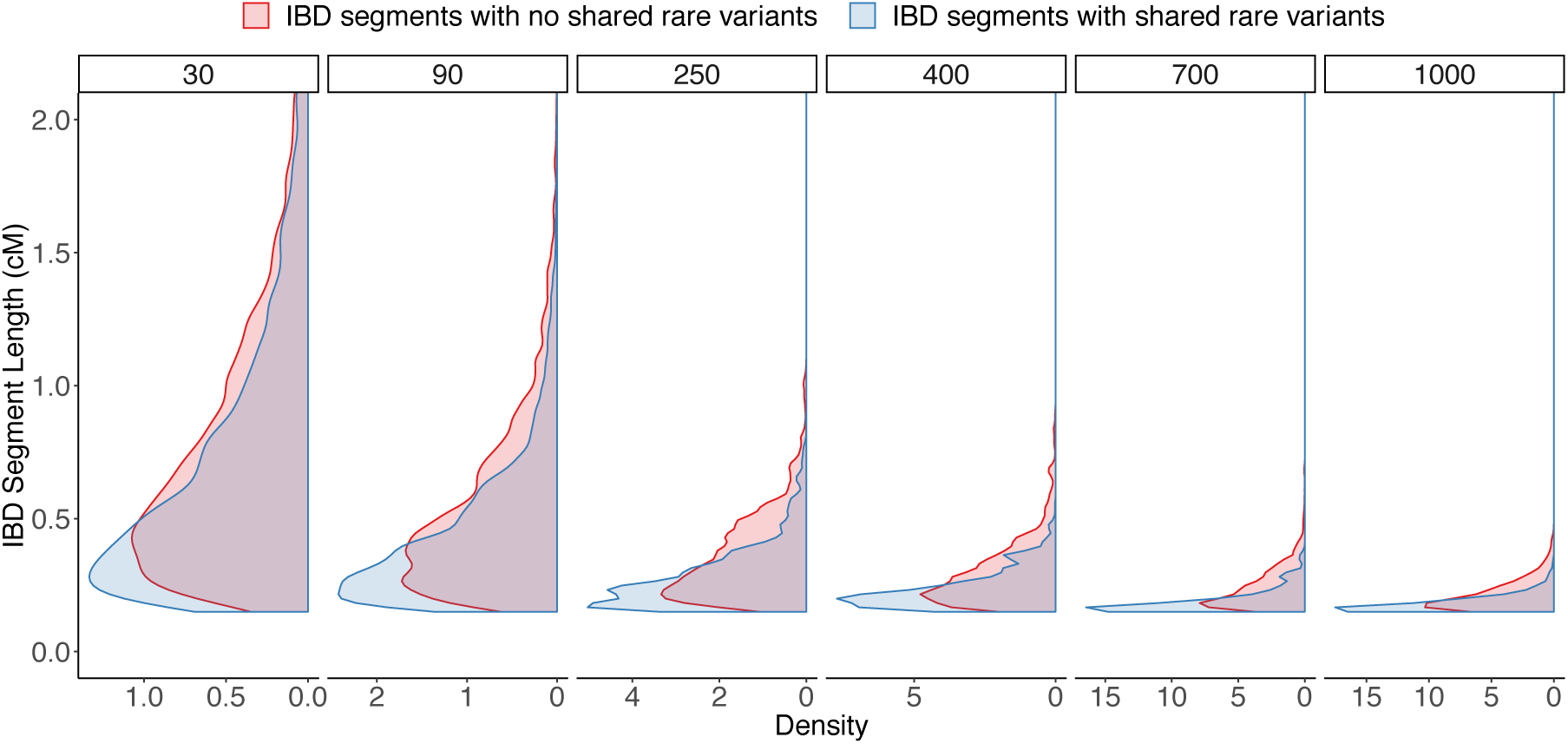
Distribution of IBD segment length grouped by presence/absence of shared rare variants. IBD segment were binned based on presence (blue) or absence (red) of shared rare variants. The distributions of segment length were compared across the two bins. Analysis was repeated for varying divergence times from left to right ranging from recent (30 generations ago) to older (1000 generations ago) divergence between POP_2_ and POP_3_ (i.e. T_POP2-POP3_).

For all simulations, we see the density of segment lengths skewed towards shorter segments. However, when we focus on short IBD segments (< 2cM), we observe a significant shift in the mode of the distribution wherein IBD segments with shared rare variants are shorter than segments with no presence of shared rare variants. In addition, as we transition from a recent divergence time (i.e. ∼750ya) to an older divergence time (i.e. ∼25,000ya), we observe the expected decrease in the range of IBD segment lengths and prominent difference between the two categories. Hence, we conclude that shared rare variants would better tag short IBD segments and allow us to assess older divergence time between populations.

### Shared rare variants within short IBD segments improve odds of IBD vs IBS

Next, we binned the IBD segments based on the original log-odds (LOD) from Refined IBD and the enriched rvIBD log-odds (rvLOD) into those that satisfy one of LOD > 3 or rvLOD > 3 or both conditions. Since the log-odds is the base 10 logarithm scaled ratio of the likelihood of IBD to the likelihood of non-IBD, a threshold of 3 represents a 1000-fold difference increase in the likelihood of IBD. We then compared the distribution of IBD segment length between the three sets (Figure 3; <u>Supplementary Figure S3</u>). Again, we observe a significant shift (skewed towards shorter segments) in the distribution of segment lengths for segments satisfying the rvLOD > 3 criterion than those that satisfy both or only the LOD > 3 criterion. We also see a decrease in the expected IBD segment lengths as expected and prominent differences between the three categories as we transition from a recent to older divergence time between the two populations.

**Figure 3:**
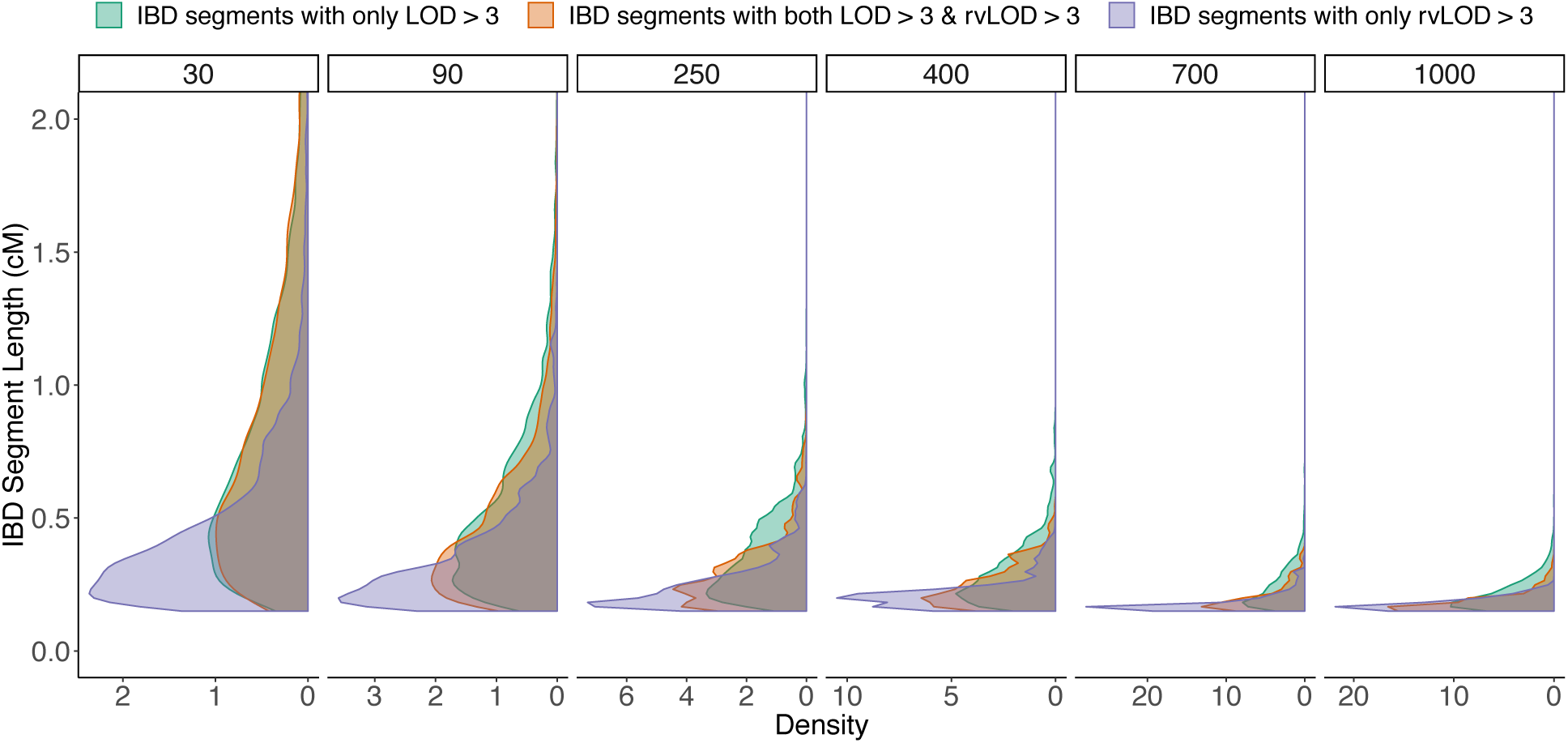
Distribution of IBD segment length categorized by LOD and rvLOD thresholds. IBD segment were binned based on satisfying one or both thresholds of LOD > 3 and/or rvLOD > 3. The distributions of segment length were compared across the three bins. Analysis was repeated for varying divergence times from left to right ranging from recent (30 generations ago) to older (1000 generations ago) divergence between POP_2_ and POP_3_ (i.e. T_POP2-POP3_).

### Short IBD segments leveraged by shared rare variants identify older relatedness

We compared the distribution of TMRCA estimates for the same three categories segregated based on LOD and rvLOD scores, as described above (Figure 4; <u>Supplementary Figure S3</u>). Since IBD segments are broken down over generations, the expected TMRCA estimates should be older as the IBD segment length decreases.

**Figure 4:**
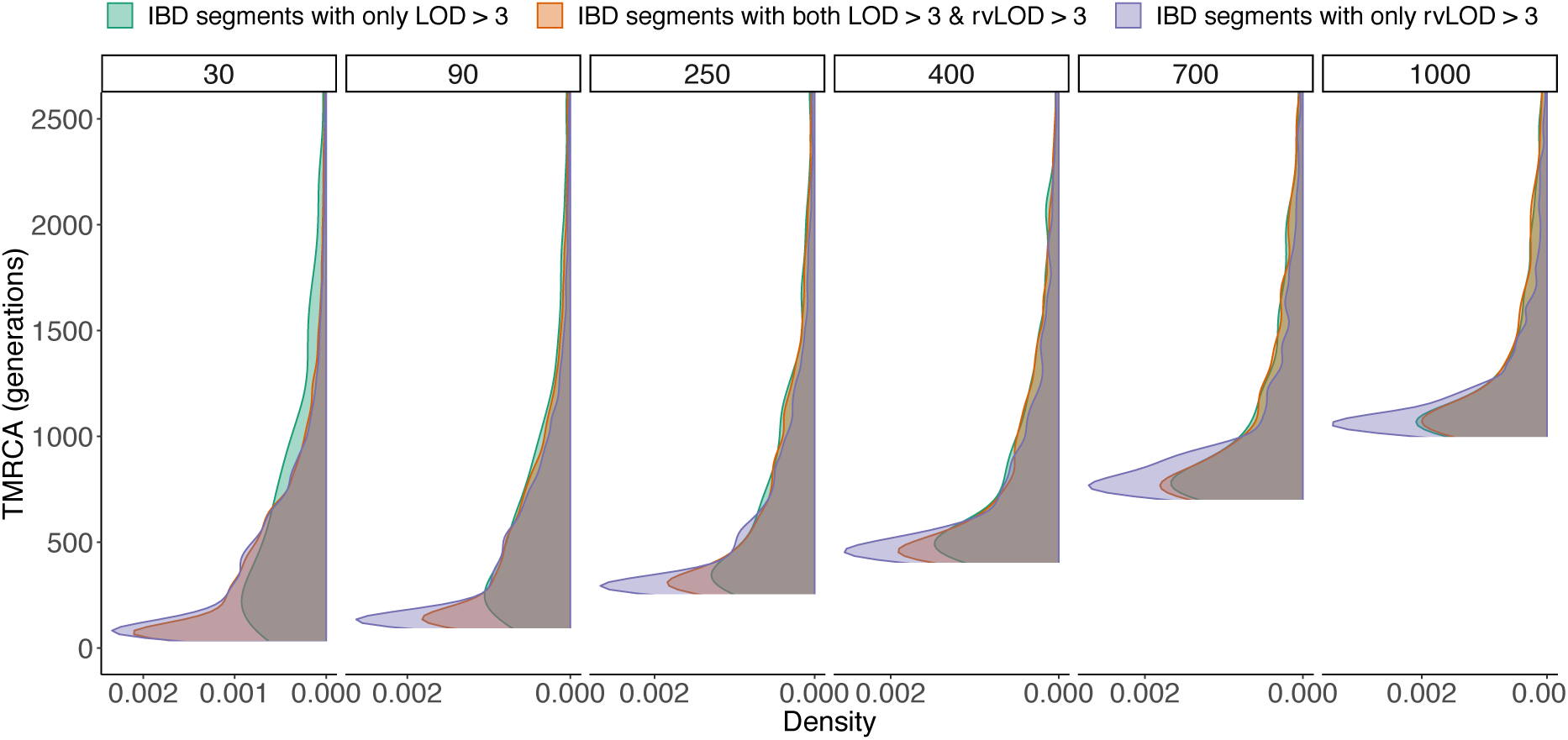
Distribution of TMRCA estimates categorized by LOD and rvLOD thresholds. The time to most recent common ancestor (TMRCA) was estimated for each IBD segment using the ancestry trees generated by *msprime*. IBD segment were binned based on satisfying one or both thresholds of LOD > 3 and/or rvLOD > 3. The distributions of TMRCA estimates were compared across the three bins. Analysis was repeated for varying divergence times from left to right ranging from recent (30 generations ago) to older (1000 generations ago) divergence between POP_2_ and POP_3_ (i.e. T_POP2-POP3_).

For each of the simulated datasets, we do observe that the distribution of TMRCA estimates agrees with the divergence time utilized to simulate the dataset. While the distribution of the TMRCA estimates for segments with only LOD > 3 peaks close to the divergence time, we see that the distribution of the TMRCA estimates for segments with rvLOD > 3 extend to estimates even older than the divergence time detecting inheritance of IBD segments from common ancestors before the separation of the two populations.

This demonstrates that the new rvIBD metric is not only able to enrich the detection of short IBD segments but also identify IBD segments corresponding to older relatedness as simulated by the different divergence times.

### Empirical dataset

In order to assess the application of the rvIBD metric to detect distant relatedness, we used the high-coverage whole genome sequencing data from freeze3 dataset of the Trans-Omics for Precision Medicine (TOPMed) Project (Taliun et al. 2019) that includes ∼1,100 individuals from the Old Order Amish (OOA) cohort in Lancaster County, Pennsylvania. The OOA population immigrated to the Colonies from Western Europe in the early 1700’s to escape religious persecution (Cross 1976). There are now over 38,000 Amish individuals in the Lancaster County, nearly all of whom can trace their ancestry back 12-14 generations to approximately 200 founders (McKusick 1978; Beiler 1988; Lee et al. 2010). The complex multi-generational pedigree allows us to assess genetic sharing between 1^st^ and 13^th^ order related individual pairs. The inclusion of both closely-related (less than 3^rd^ order) and higher order relatedness enables us to assess the accuracy of the rvIBD metric for different orders of relatedness.

Using markers with a minimum minor allele frequency of 5%, we identified IBD segments using default parameters except for lowering the length threshold to 0.5cM and the log-odds threshold to 1. For the assessment of the rvIBD metric, we filtered the possible ∼600,000 pairs of individuals to retain only individual-pairs with non-consanguineous common ancestors as inferred from the pedigree structure. Each IBD segment detected using *Refined IBD* was leveraged for the presence of shared rare variants (minor allele frequency < 1%) to compute the number of shared rare variants and the enriched rvIBD log-odds (rvLOD) score.

For each pair of individuals, we filtered the set of IBD segments based on the different segment length thresholds, thresholds for LOD score and thresholds for rvLOD scores. The observed IBD proportion for each pair was computed as the ratio of the cumulative length of IBD segments (in cM) retained after the filtering process to the length of the genome (in cM). For each pair of individuals, we also estimated the order of relatedness inferred from the number of generations to their most recent common ancestor in the pedigree. The expected IBD proportion was computed as described by Manichaikul et. al (2010). Further, we estimated the precision of detection of short IBD segments by comparing IBD segments shared between distant cousins to the IBD segments shared between one of the cousins and the parents/grandparents of the other cousin, when available. The precision was calculated for different segment bins and assessed for different orders of relatedness.

### Cryptic genetic relatedness in the Older Order Amish cohort

Overall, in the OOA dataset, we illustrate large estimates of IBD sharing between individuals expected due to the cohort originating from a founder population. We compared the observed IBD proportions to the expected IBD proportions after using different sets of thresholds for IBD segment length, original LOD score and rvLOD score based on the rvIBD algorithm (Figure 5; <u>Supplementary Figure S4</u>).

**Figure 5:**
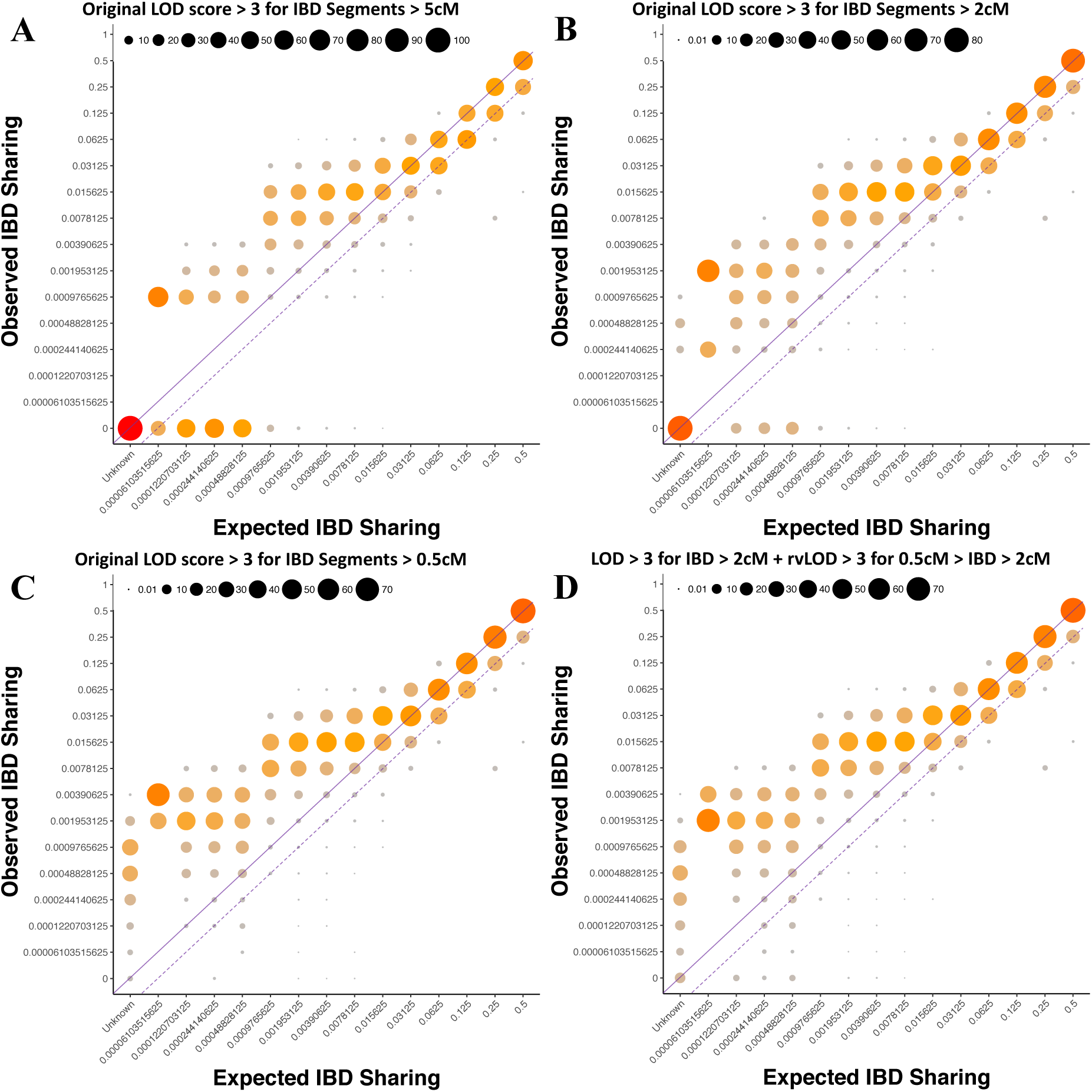
Performance of relatedness estimation using different sets of IBD segment thresholds. Comparisons of observed IBD proportion using genetic data was made to the expected IBD proportion estimated from the pedigree for different degrees of relatedness using (A) IBD segments > 5cM with LOD > 3, (B) IBD segments > 2cM with LOD > 3, (C) IBD segments > 0.5cM with LOD > 3, and (D) IBD segments > 2cM with LOD > 3 and IBD segments < 2cM with rvLOD > 3. Area of the circles indicates the percentage of individual pairs with specific proportions of observed IBD sharing segregated by the estimated degrees of relationship. The solid line represents bins where the observed IBD proportions were comparable to the expected IBD proportions while the dotted line represents a lower threshold used to estimate the proportion of individual pairs with detectable relatedness based on genetic IBD sharing illustrated in Supplementary Figure S4.

When using a minimum LOD score of 3 and minimum segment length of 5cM (Figure 5A), we observe more than the expected IBD sharing in 90% of pairs for relatedness order ranging from 1^st^ to 9^th^ degree relatives with no IBD sharing estimated in the seemingly unrelated individuals. When we decrease the segment length threshold to 2cM (Figure 5B), we observe similar results with IBD sharing among some unrelated individuals while a lenient length threshold of 0.5cM (Figure 5C) results in IBD sharing detected in over 90% pairs across all orders of relatedness as well as over 80% of the seemingly unrelated pairs. This sizeable increase in observed IBD sharing proportions could be caused by the introduction of false positive IBD segments ranging from 0.5cM to 2cM in length.

In order to reduce these false positives, we implemented a composite threshold of LOD > 3 for segment lengths greater than 2cM and rvLOD > 3 for segment lengths less than 2cM (Figure 5D) and observed IBD sharing in 90% of pairs for relatedness order ranging from 1^st^ to 9^th^ degree relatives with greater than 10% increase in the detection rate of 5^th^ cousins (10^th^ order) and more distant relatedness. IBD sharing in the seemingly unrelated individuals rose from a single individual sharing IBD with other individuals. Detection of possible cryptic relatedness among the seemingly unrelated individuals with potential common ancestors older than 14 generations ago cannot be assessed due to limited pedigree information and lack of knowledge regarding the relatedness among founders of the Old-Order Amish community.

Lastly, we assessed the precision of detecting short IBD segments using the original LOD estimates and the rvLOD estimates. For each degree of relationships ranging from 4^th^ to 13^th^ order relatedness, we segregated the IBD segments by length into 7 groups. For a given IBD segment between a pair of cousins, we assessed if an overlapping segment was also detected between one of the cousins and the parents or grandparents of the other cousin under the premise that the genomic region was inherited from one of the two parents or one of the 4 grandparents, when present in the sequenced OOA cohort. We considered the IBD to be true if overlapping IBD segments were detected and false if overlapping segments were not found. The precision of detecting IBD segments of a particular length was computed based on the count of true IBD and false IBD segments. The count of false IBD segments was scaled when full pedigree information was unavailable. We repeated the analysis for the different thresholds defined in Figure 5. We show an improvement in the precision of detecting short segments (< 2cM) detected using the rvIBD metric over the original IBD callset (<u>Supplementary Figure S5</u>). The precision of shorter segments based on rvIBD is comparable to the precision of longer segments using the original IBD callset which are already known to have higher accuracy as demonstrated by previous work in the field (Browning and Browning 2010; Genovese et al. 2010; Browning and Browning 2012; Browning and Browning 2013b).

## Discussion

Here we have described an approach, rvIBD, for leveraging the presence of shared rare-variants between two individuals to better assess their genetic relatedness. Genetic relatedness is a fundamental concept in population genetics and genealogical studies and traditionally described as the probability of sharing alleles that are identical-by-descent from common ancestors in a pedigree. Identity-by-descent is the phenomenon whereby two individuals inherit a genomic region from a common ancestor without an intervening detectable recombination event (Weir et al. 2006). With the onset of dense genotyping and next-generation sequencing, genetic relatedness has been computed directly from single nucleotide polymorphism (SNP) data using maximum likelihood estimates (Milligan 2003). Using averages of single SNP-based estimates, matrices of genetic relatedness have been computed and incorporated in to genome wide association studies (GWAS) (Forni et al. 2011; Yang et al. 2011; Yang et al. 2013; Chang et al. 2015). However, single SNP-based estimates do not take into account linkage disequilibrium (LD), discard SNPs based on minor allele frequency thresholds, and ignore the length of shared genomic regions between two individuals.

More recently, methods detecting haplotype sharing have been developed (Gusev et al. 2009; Browning and Browning 2010; Browning and Browning 2013b) and utilize more sophisticated methods involving hidden Markov models to differentiate IBD from IBS, which have been shown to perform better (Ramstetter et al. 2017). We specifically focus on the detection of short IBD segments (<2cM) that have been previously unreliable and are more indicative of older co-ancestry between individuals. We demonstrated through our simulations of populations with varying divergence times, the capability of rare variants to better tag short IBD segments over common variants and subsequently improve the assessment of IBD vs IBS.

In the last decade, there has been an increasing focus on the existence of cryptic relatedness in both isolated and cosmopolitan genetic samples (Voight and Pritchard 2005; Pemberton et al. 2010; Henn et al. 2012; Fedorova et al. 2016; Staples et al. 2018). Cryptic relatedness refers to the presence of relatives in a sample of seemingly unrelated individuals. Voight and Pritchard showed that cryptic relatedness has a significant inflation of association p-values from genetic association studies involving small and isolated populations resulting in false positives. First, Pemberton and colleagues were able to infer previously unidentified relationships in phase III of the HapMap Project (Pemberton et al. 2010). Later, Henn and colleagues detected thousands of higher-order relationships in a heterogeneous European population (Henn et al. 2012). Recently, Staples and colleagues highlighted the prevalence of cryptic relatedness in large healthcare studies (Staples et al. 2018).

Through our analyses of simulated data, we show the ability of detecting short IBD segments by leveraging shared rare variants. We demonstrated that these short segments when enriched with shared rare variants are indicative of older relatedness between individuals from populations that diverged many thousands of years in the past. Furthermore, using the empirical dataset from the multi-generational Older Order Amish cohort, we illustrate the improvement in the detection of higher-order relationships greater than fifth-cousins after the inclusion of the rvIBD metric for short IBD segments. Using the short segments leveraged by shared rare variants, we were also able to identify genetic relatedness between the seemingly unrelated individuals which will need further investigation. If left undetected, these higher-order relationships contribute to the prevalence of confounding due to cryptic relatedness in large healthcare studies.

Recent large-scale human sequencing projects have identified tens of millions of variants (1000 Genomes Project Consortium et al. 2012; 1000 Genomes Project Consortium et al. 2015; Sudlow et al. 2015; Dewey et al. 2016; Gaziano et al. 2016; Taliun et al. 2019), most of which are rare variants (minor allele frequency < 1%). Common variants are often shared by chance with no underlying IBD segments. Based on the infinite-site model and the low rates of mutation, the possibility of a rare variant shared between two individuals without common ancestry is less likely. Hence, the sharing of rare variants is more indicative of co-ancestry than the sharing of common variants. Recent explosive population growth has resulted in a deluge of rare genetic variants due to the increased mutational capacity of recent human populations (Coventry et al. 2010; Keinan and Clark 2012; Fu et al. 2013). Fu and colleagues showed that a majority of the variants are predicted to have arisen in the last 10,000 to 50,000 years (Fu et al. 2013). Work by O’Connor and colleagues further demonstrated that rare variants have more information content than common variants when making inferences of fine-scale population structure since they can accurately detect recent demographic changes in the last 25,000 years (O’Connor et al. 2015). Other works focused on rare variants also showed the ability to identify distant relatedness within and between cohorts (Hochreiter 2013; Mathieson and McVean 2014; Al-Khudhair et al. 2015).

Overall, we show that leveraging shared rare variants, which were previously ignored, allows for the detection of older co-ancestry. Detection of older co-ancestry will help identify distant relatedness between populations and will provide new insights in to population structure using rare variants. In addition, detection of older co-ancestry will also help identify cryptic relatedness in large-scale studies involving cohorts ranging from ten to hundred thousand individuals which if left unidentified could lead to spurious results in downstream association analyses. Indirectly, the detection of shorter IBD segments would augment other applications for IBD segments involving the estimation of population demographic parameters, inferring fine-scale population structure, detecting migratory events, IBD mapping and forensic science (Palamara et al. 2012; Browning and Browning 2012; Thompson 2013; Palamara and Pe’er 2013; Palamara et al. 2015).

## Acknowledgements

Whole genome sequencing (WGS) for the Trans-Omics in Precision Medicine (TOPMed) program was supported by the National Heart, Lung and Blood Institute (NHLBI). WGS for “NHLBI TOPMed: Genetics of Cardiometabolic Health in the Amish” (phs000956.v1.p1) was performed at Broad Institute of MIT and Harvard (3R01HL121007-01S1). Centralized read mapping and genotype calling, along with variant quality metrics and filtering were provided by the TOPMed Informatics Research Center (3R01HL-117626-02S1; contract HHSN268201800002I). Phenotype harmonization, data management, sample-identity QC, and general study coordination were provided by the TOPMed Data Coordinating Center (3R01HL-120393-02S1; contract HHSN268201800001I). We gratefully acknowledge the studies and participants who provided biological samples and data for TOPMed. A full list of TOPMed collaborators can be found at https://www.nhlbiwgs.org/topmed-banner-authorship. The TOPMed component of the Amish Research Program was supported by NIH grants R01 HL121007, U01 HL072515, and R01 AG18728.

