## Supplementary Figures for "Rare variant enriched identity-by-descent enables the detection of distant relatedness and older divergence between populations"

**Shetty AC, et. al.**

### Supplementary Figures

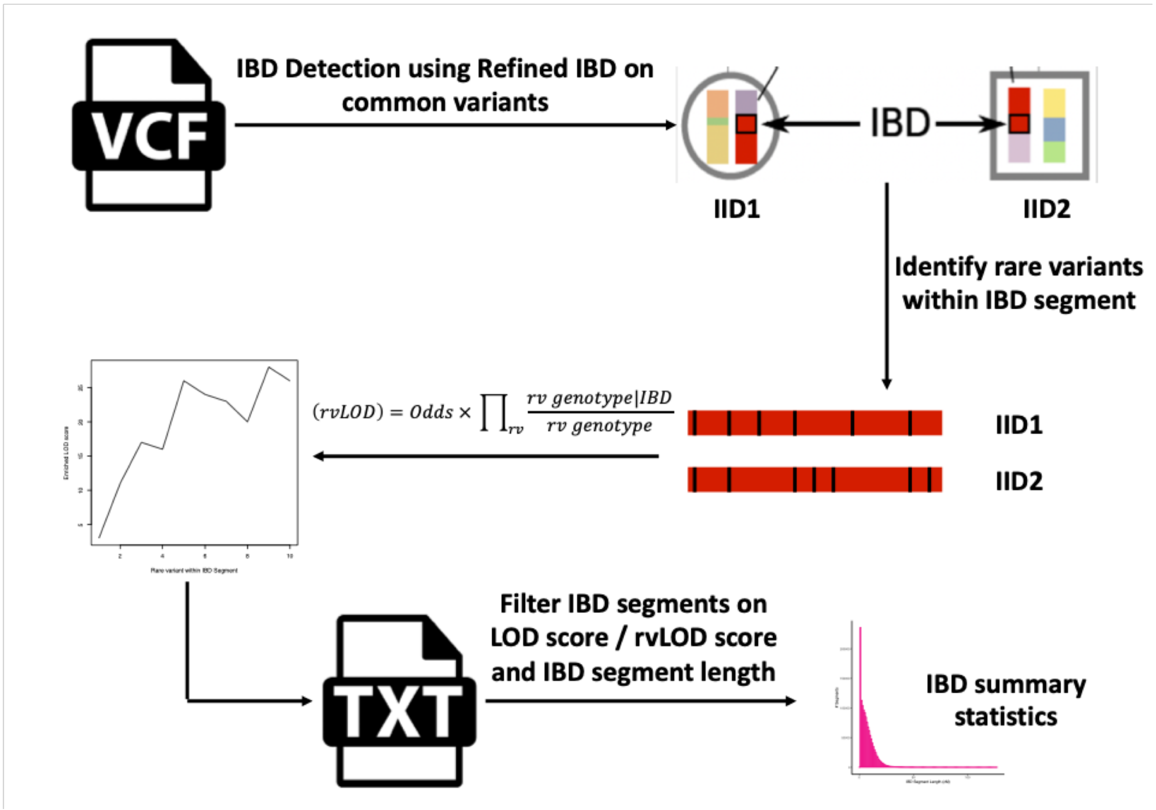


**Figure S1: Schematic of the rare variant enriched IBD (rvIBD) algorithm.**

The rare variant IBD (rvIBD) algorithm involves three steps, namely (1) the detection of IBD segments between individual pairs using Refined IBD on a set of common variants (MAF > 5%), (2) the identification of rare variants within the endpoints of the IBD segment, and (3) the computation of the rvIBD metric using a Bayesian approach to enrich the odds of IBD vs IBS by leveraging the sharing of rare variants within the IBD segment. Finally, the IBD segments can be filtered based on different threshold sets for the original log-odds (LOD) score, the enriched log-odds (rvLOD), and the length of the IBD segment.

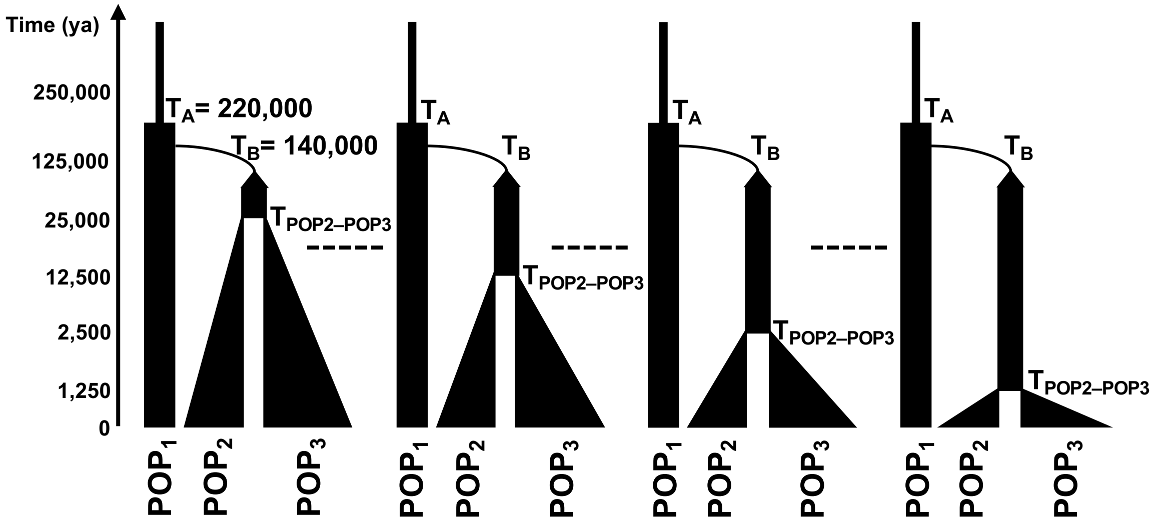


| | |
|---|---|
| $T_A$ | Ancestral population size of 12,300 about 220,000 ya |
| $T_B$ | Out-of-$POP_1$ reduction to 2,100 occurring 140,000 ya |
| $T_{POP2\text{-}POP3}$ | Divergence between $POP_2$ and $POP_3$ of size 1000 and 510 respectively ranging from 125 – 25,000 years before present |

**Figure S2: Population models used for simulated datasets.**

We fit the population model proposed by Gutenkunst et. al. (Gutenkunst et al. 2009) and simulated multiple datasets after varying the divergence time between $POP_2$ and $POP_3$ from 125 years before present to 25,000 years before present in order to simulate recent to distant relatedness between individuals from $POP_2$ and $POP_3$.

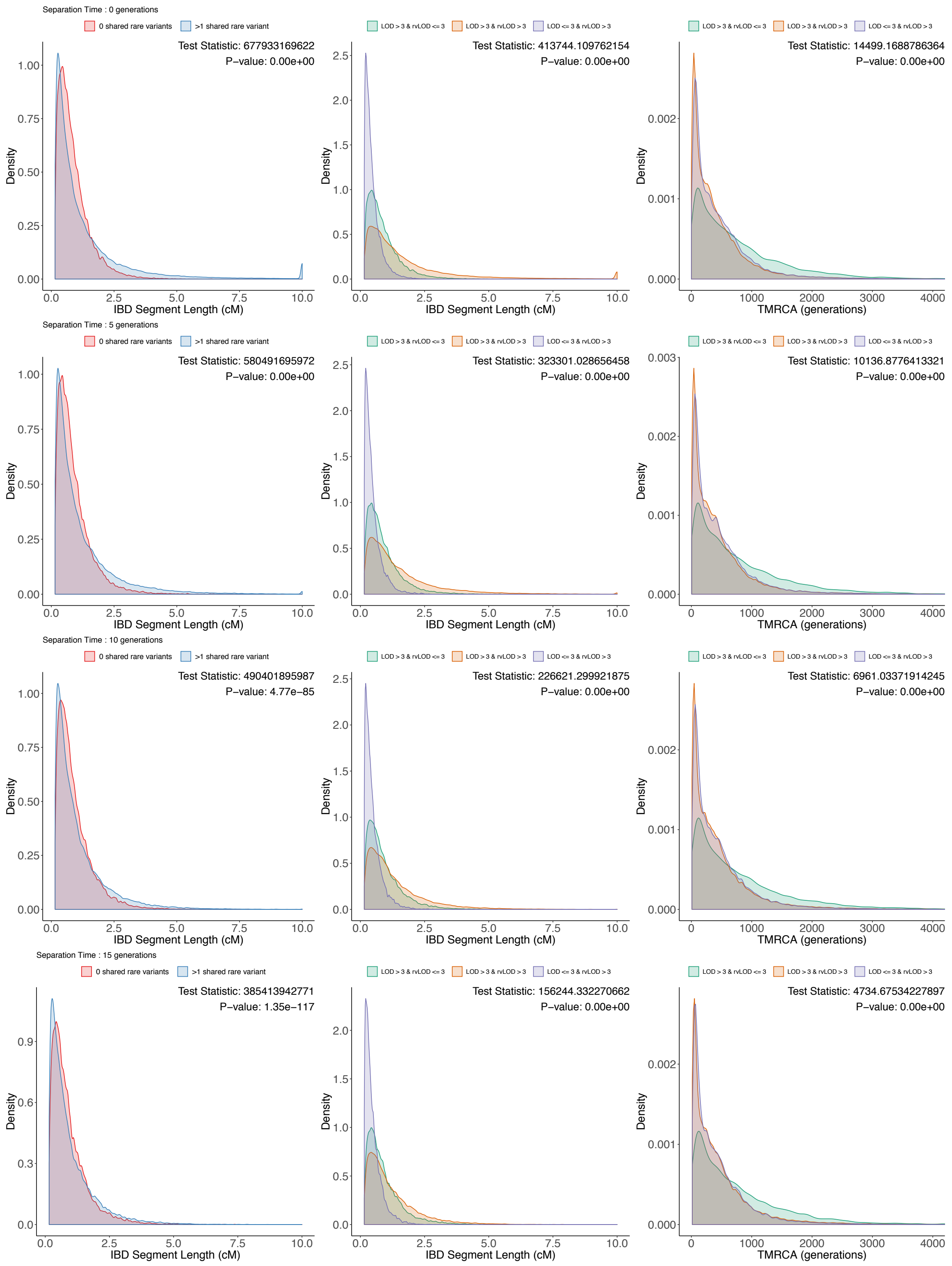
Separation Time : 0 generations
0 shared rare variants
>1 shared rare variant
Test Statistic: 677933169622
P−value: 0.00e+00
LOD > 3 & rvLOD <= 3
LOD > 3 & rvLOD > 3
LOD <= 3 & rvLOD > 3
Test Statistic: 413744.109762154
P−value: 0.00e+00
Test Statistic: 14499.1688786364
P−value: 0.00e+00
Separation Time : 5 generations
Test Statistic: 580491695972
P−value: 0.00e+00
Test Statistic: 323301.028656458
P−value: 0.00e+00
Test Statistic: 10136.8776413321
P−value: 0.00e+00
Separation Time : 10 generations
Test Statistic: 490401895987
P−value: 4.77e−85
Test Statistic: 226621.299921875
P−value: 0.00e+00
Test Statistic: 6961.03371914245
P−value: 0.00e+00
Separation Time : 15 generations
Test Statistic: 385413942771
P−value: 1.35e−117
Test Statistic: 156244.332270662
P−value: 0.00e+00
Test Statistic: 4734.67534227897
P−value: 0.00e+00
Density
IBD Segment Length (cM)
TMRCA (generations)

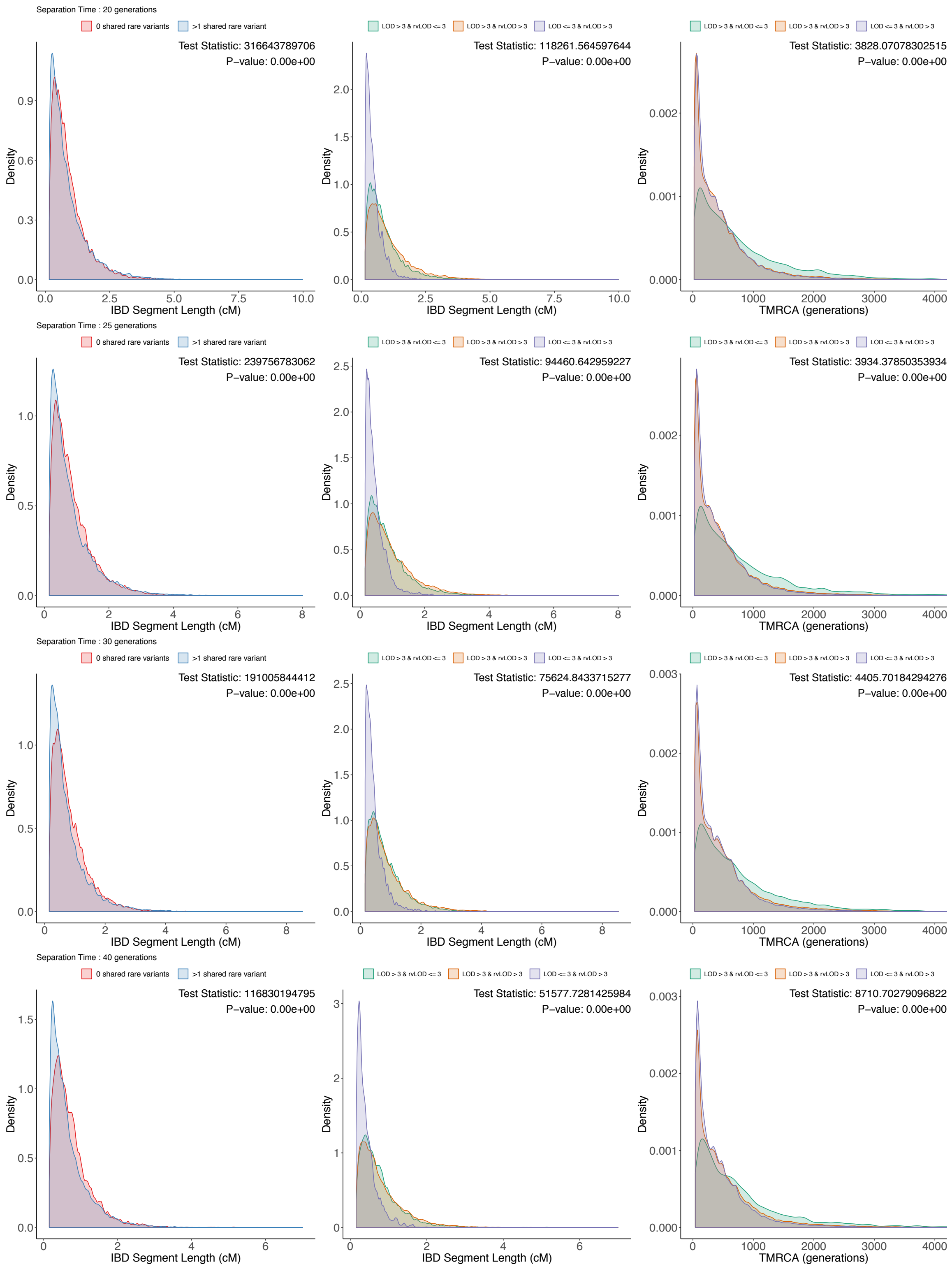

Separation Time : 20 generations
0 shared rare variants
>1 shared rare variant
Test Statistic: 316643789706
P−value: 0.00e+00
LOD > 3 & rvLOD <= 3
LOD > 3 & rvLOD > 3
LOD <= 3 & rvLOD > 3
Test Statistic: 118261.564597644
P−value: 0.00e+00
Test Statistic: 3828.07078302515
P−value: 0.00e+00
Separation Time : 25 generations
Test Statistic: 239756783062
P−value: 0.00e+00
Test Statistic: 94460.642959227
P−value: 0.00e+00
Test Statistic: 3934.37850353934
P−value: 0.00e+00
Separation Time : 30 generations
Test Statistic: 191005844412
P−value: 0.00e+00
Test Statistic: 75624.8433715277
P−value: 0.00e+00
Test Statistic: 4405.70184294276
P−value: 0.00e+00
Separation Time : 40 generations
Test Statistic: 116830194795
P−value: 0.00e+00
Test Statistic: 51577.7281425984
P−value: 0.00e+00
Test Statistic: 8710.70279096822
P−value: 0.00e+00
Density
IBD Segment Length (cM)
TMRCA (generations)

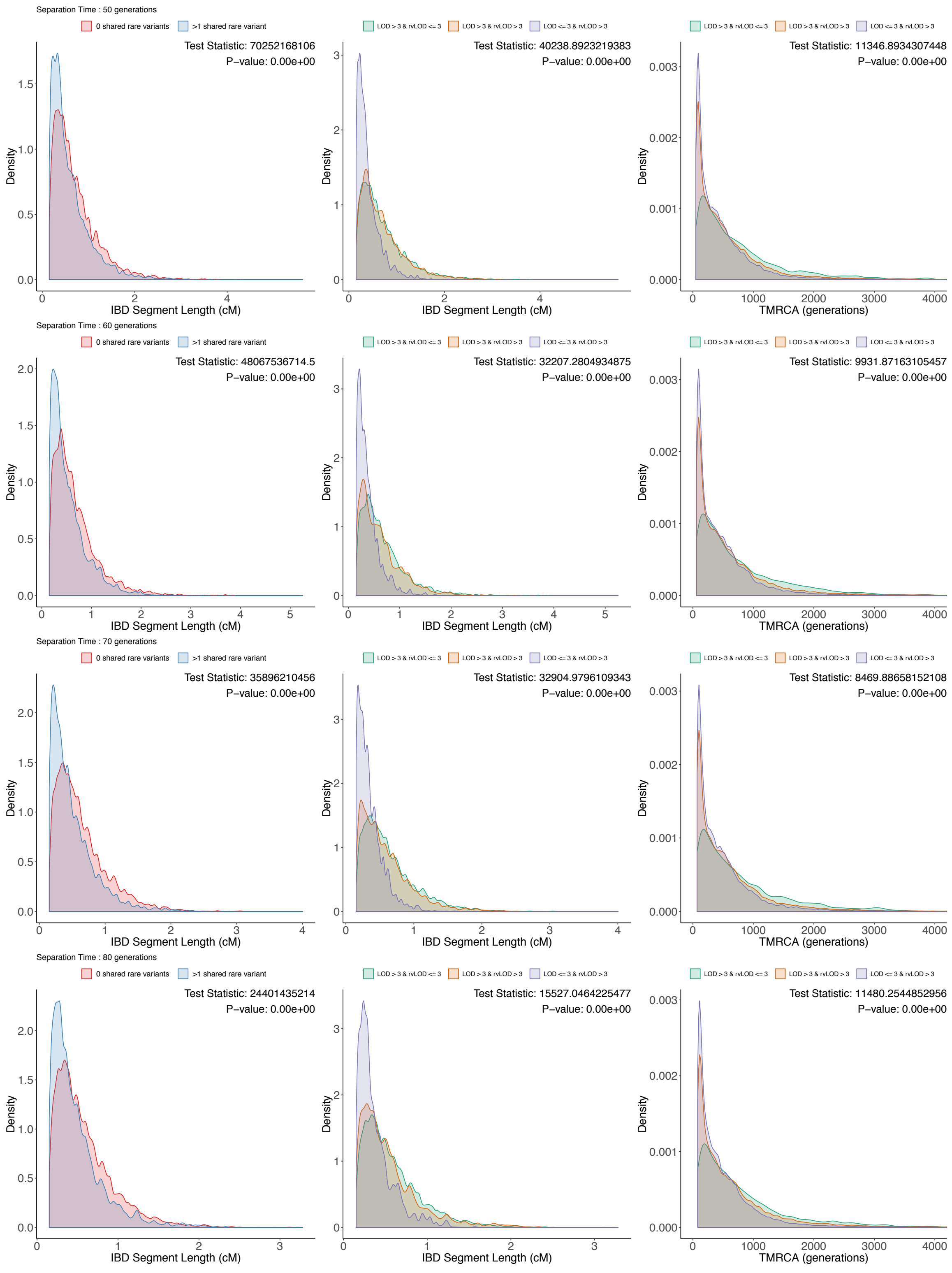

Separation Time : 50 generations
0 shared rare variants
>1 shared rare variant
Test Statistic: 70252168106
P−value: 0.00e+00
LOD > 3 & rvLOD <= 3
LOD > 3 & rvLOD > 3
LOD <= 3 & rvLOD > 3
Test Statistic: 40238.8923219383
P−value: 0.00e+00
Test Statistic: 11346.8934307448
P−value: 0.00e+00
Separation Time : 60 generations
Test Statistic: 48067536714.5
P−value: 0.00e+00
Test Statistic: 32207.2804934875
P−value: 0.00e+00
Test Statistic: 9931.87163105457
P−value: 0.00e+00
Separation Time : 70 generations
Test Statistic: 35896210456
P−value: 0.00e+00
Test Statistic: 32904.9796109343
P−value: 0.00e+00
Test Statistic: 8469.88658152108
P−value: 0.00e+00
Separation Time : 80 generations
Test Statistic: 24401435214
P−value: 0.00e+00
Test Statistic: 15527.0464225477
P−value: 0.00e+00
Test Statistic: 11480.2544852956
P−value: 0.00e+00
Density
IBD Segment Length (cM)
TMRCA (generations)

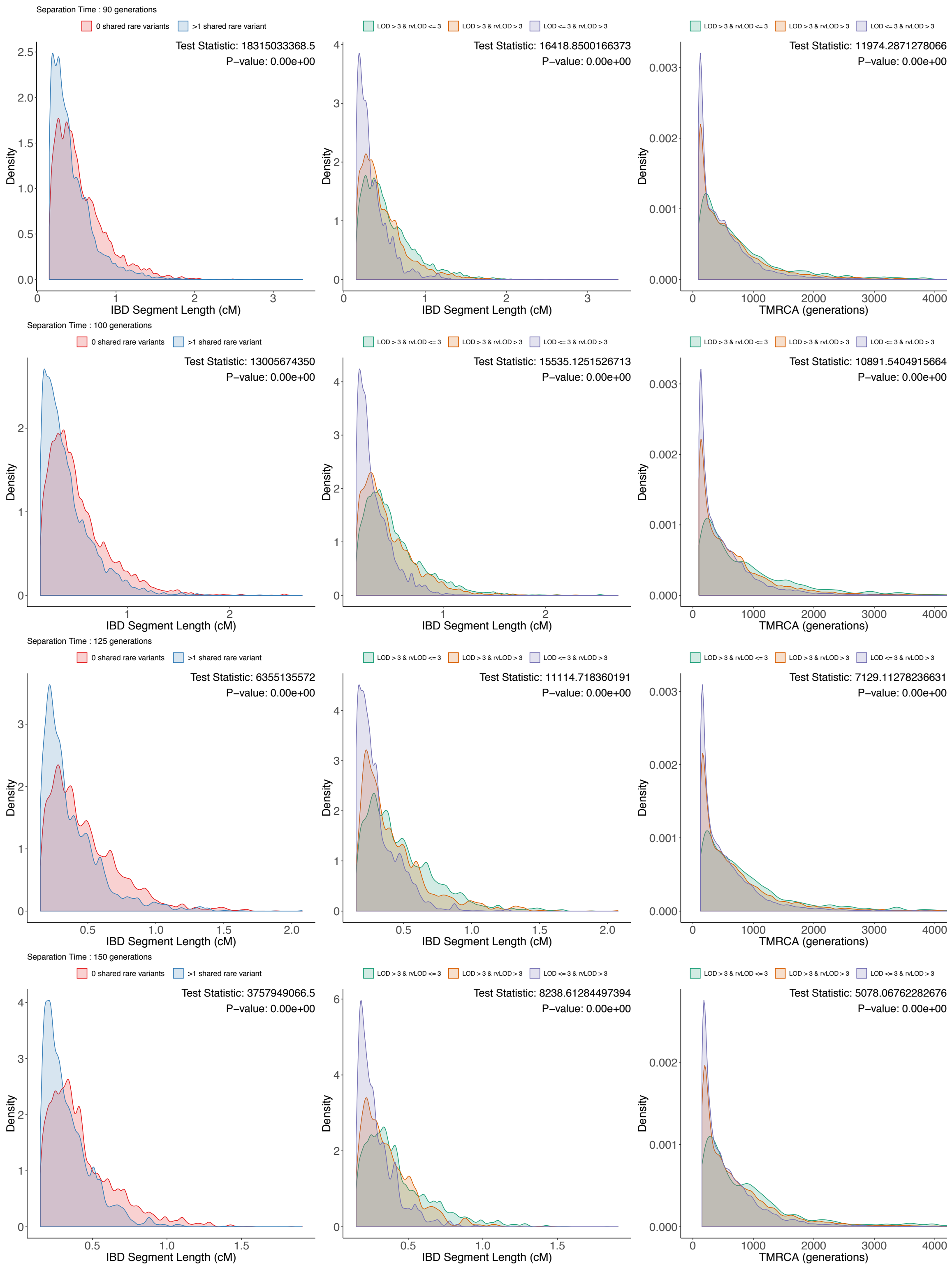

Separation Time : 90 generations
0 shared rare variants
>1 shared rare variant
Test Statistic: 18315033368.5
P−value: 0.00e+00
Density
IBD Segment Length (cM)
LOD > 3 & rvLOD <= 3
LOD > 3 & rvLOD > 3
LOD <= 3 & rvLOD > 3
Test Statistic: 16418.8500166373
P−value: 0.00e+00
Test Statistic: 11974.2871278066
P−value: 0.00e+00
TMRCA (generations)
Separation Time : 100 generations
Test Statistic: 13005674350
P−value: 0.00e+00
Test Statistic: 15535.1251526713
P−value: 0.00e+00
Test Statistic: 10891.5404915664
P−value: 0.00e+00
Separation Time : 125 generations
Test Statistic: 6355135572
P−value: 0.00e+00
Test Statistic: 11114.718360191
P−value: 0.00e+00
Test Statistic: 7129.11278236631
P−value: 0.00e+00
Separation Time : 150 generations
Test Statistic: 3757949066.5
P−value: 0.00e+00
Test Statistic: 8238.61284497394
P−value: 0.00e+00
Test Statistic: 5078.06762282676
P−value: 0.00e+00

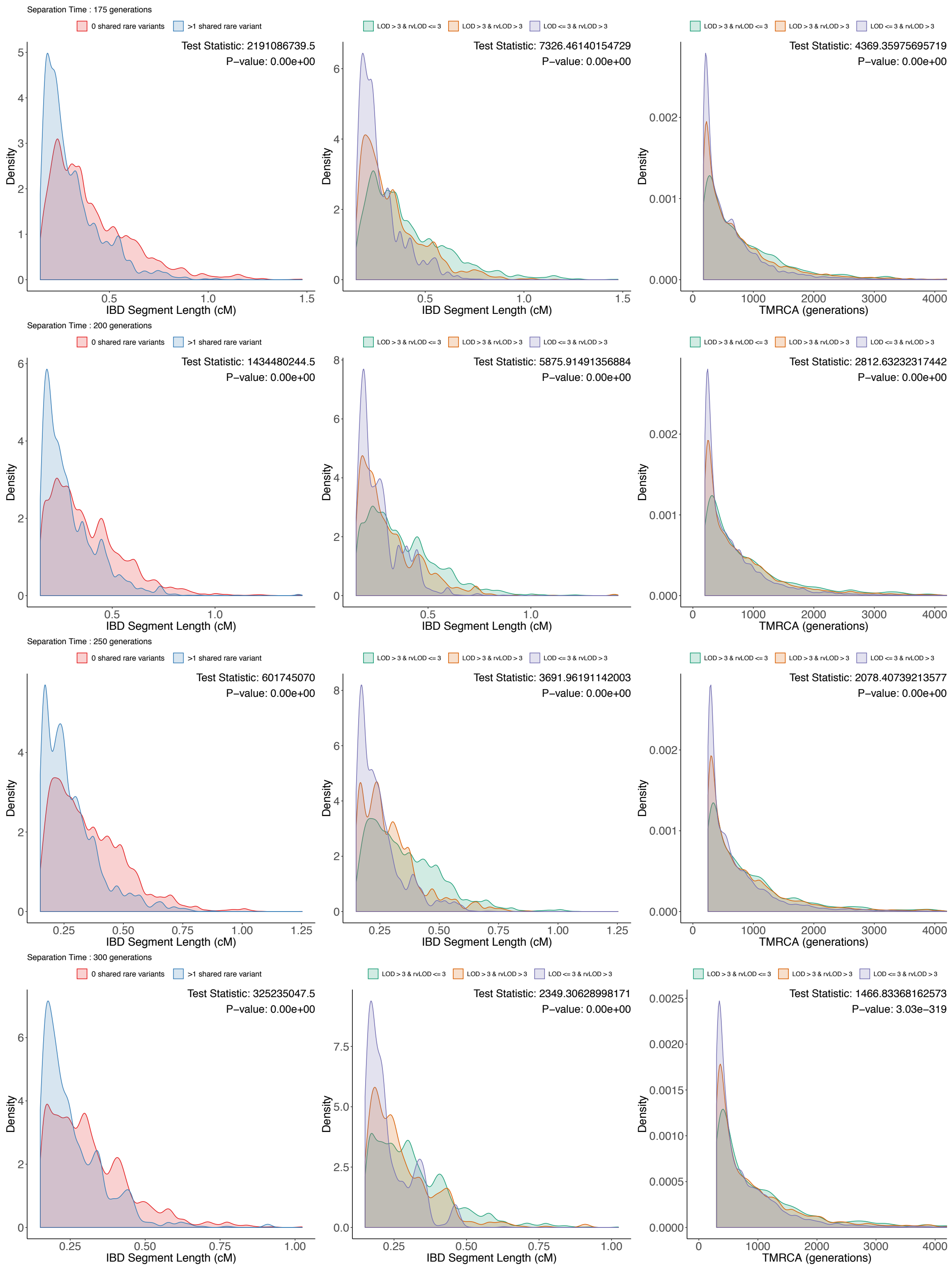
Separation Time : 175 generations
0 shared rare variants
>1 shared rare variant
Test Statistic: 2191086739.5
P−value: 0.00e+00
LOD > 3 & rvLOD <= 3
LOD > 3 & rvLOD > 3
LOD <= 3 & rvLOD > 3
Test Statistic: 7326.46140154729
P−value: 0.00e+00
Test Statistic: 4369.35975695719
P−value: 0.00e+00
Separation Time : 200 generations
Test Statistic: 1434480244.5
P−value: 0.00e+00
Test Statistic: 5875.91491356884
P−value: 0.00e+00
Test Statistic: 2812.63232317442
P−value: 0.00e+00
Separation Time : 250 generations
Test Statistic: 601745070
P−value: 0.00e+00
Test Statistic: 3691.96191142003
P−value: 0.00e+00
Test Statistic: 2078.40739213577
P−value: 0.00e+00
Separation Time : 300 generations
Test Statistic: 325235047.5
P−value: 0.00e+00
Test Statistic: 2349.30628998171
P−value: 0.00e+00
Test Statistic: 1466.83368162573
P−value: 3.03e−319
Density
IBD Segment Length (cM)
TMRCA (generations)

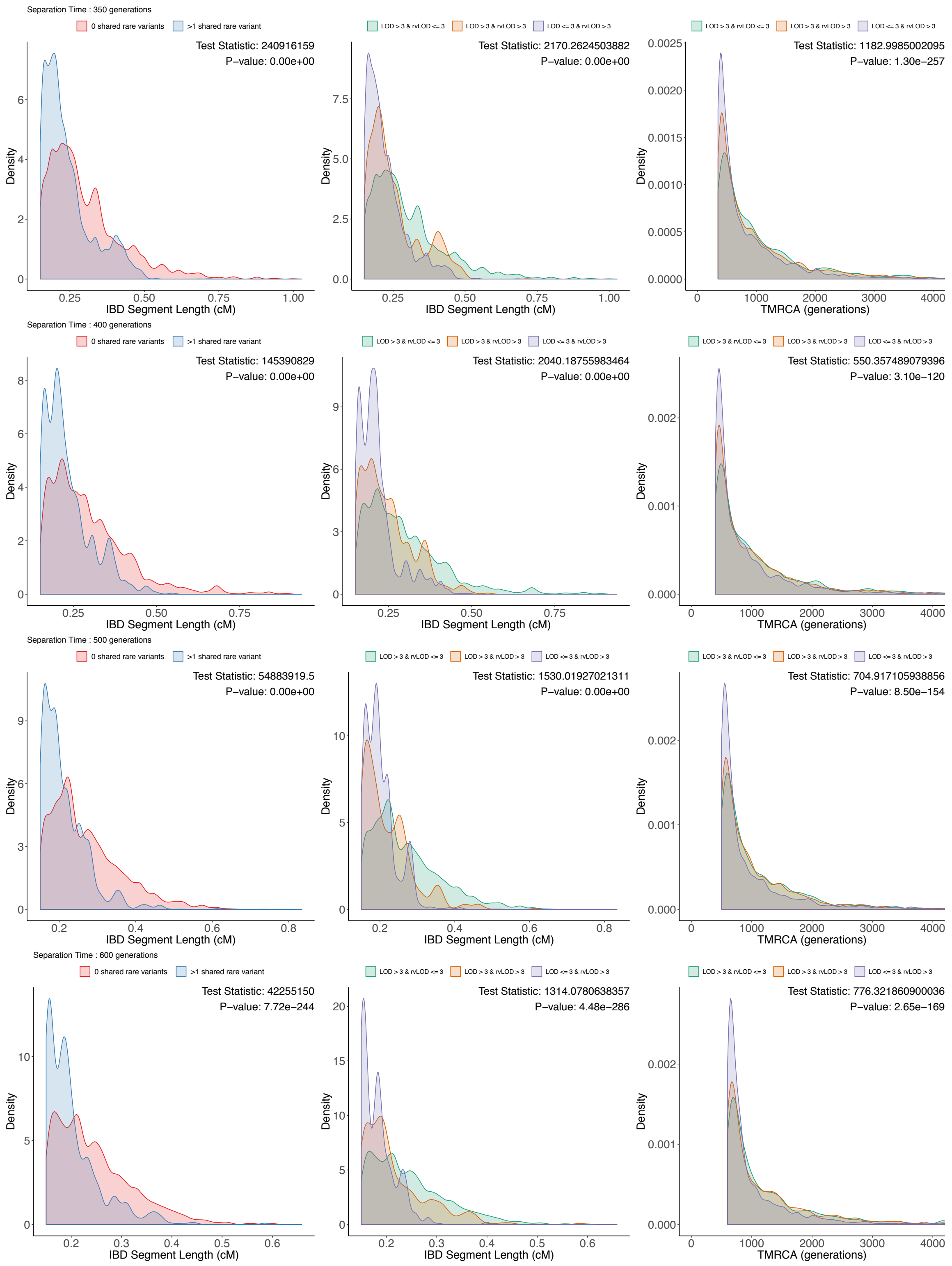
Separation Time : 350 generations
0 shared rare variants
>1 shared rare variant
Test Statistic: 240916159
P−value: 0.00e+00
Density
IBD Segment Length (cM)
LOD > 3 & rvLOD <= 3
LOD > 3 & rvLOD > 3
LOD <= 3 & rvLOD > 3
Test Statistic: 2170.2624503882
P−value: 0.00e+00
Density
IBD Segment Length (cM)
LOD > 3 & rvLOD <= 3
LOD > 3 & rvLOD > 3
LOD <= 3 & rvLOD > 3
Test Statistic: 1182.9985002095
P−value: 1.30e−257
Density
TMRCA (generations)
Separation Time : 400 generations
0 shared rare variants
>1 shared rare variant
Test Statistic: 145390829
P−value: 0.00e+00
Density
IBD Segment Length (cM)
LOD > 3 & rvLOD <= 3
LOD > 3 & rvLOD > 3
LOD <= 3 & rvLOD > 3
Test Statistic: 2040.18755983464
P−value: 0.00e+00
Density
IBD Segment Length (cM)
LOD > 3 & rvLOD <= 3
LOD > 3 & rvLOD > 3
LOD <= 3 & rvLOD > 3
Test Statistic: 550.357489079396
P−value: 3.10e−120
Density
TMRCA (generations)
Separation Time : 500 generations
0 shared rare variants
>1 shared rare variant
Test Statistic: 54883919.5
P−value: 0.00e+00
Density
IBD Segment Length (cM)
LOD > 3 & rvLOD <= 3
LOD > 3 & rvLOD > 3
LOD <= 3 & rvLOD > 3
Test Statistic: 1530.01927021311
P−value: 0.00e+00
Density
IBD Segment Length (cM)
LOD > 3 & rvLOD <= 3
LOD > 3 & rvLOD > 3
LOD <= 3 & rvLOD > 3
Test Statistic: 704.917105938856
P−value: 8.50e−154
Density
TMRCA (generations)
Separation Time : 600 generations
0 shared rare variants
>1 shared rare variant
Test Statistic: 42255150
P−value: 7.72e−244
Density
IBD Segment Length (cM)
LOD > 3 & rvLOD <= 3
LOD > 3 & rvLOD > 3
LOD <= 3 & rvLOD > 3
Test Statistic: 1314.0780638357
P−value: 4.48e−286
Density
IBD Segment Length (cM)
LOD > 3 & rvLOD <= 3
LOD > 3 & rvLOD > 3
LOD <= 3 & rvLOD > 3
Test Statistic: 776.321860900036
P−value: 2.65e−169
Density
TMRCA (generations)

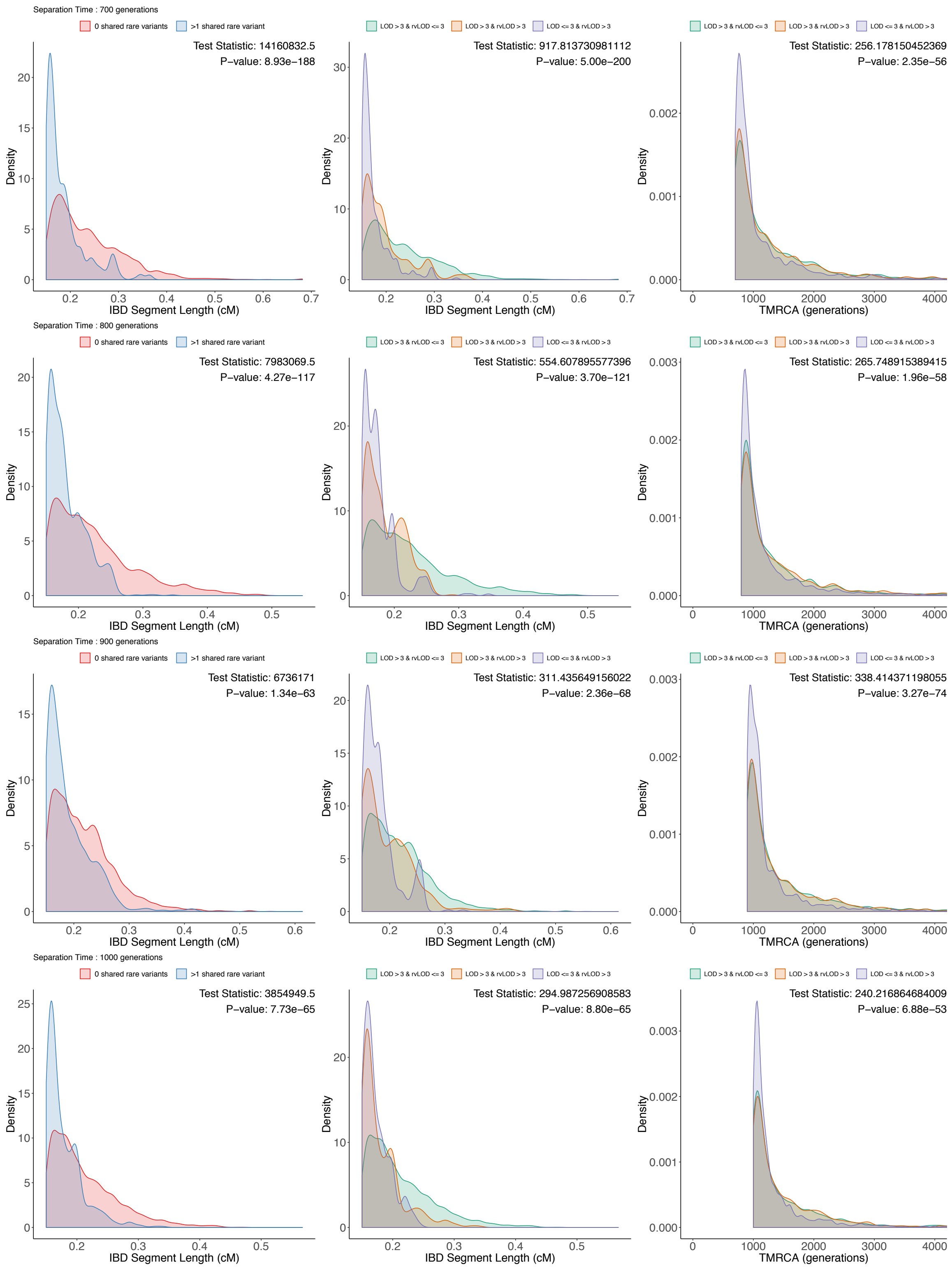


**Figure S3: Distribution of IBD segment lengths and their respective TMRCA for simulated datasets.**

For each simulated dataset (each row), we illustrate the distribution of IBD segment length binned by rare-variant sharing (column 1) and categorized based on original and rare variant LOD score (column 2).

We also illustrate the distribution of TMRCA (column 3) for the IBD segments categorized based on original and rare variant LOD score. Differences were tested using Wilcox rank-sum or Kruskal-Wallis tests.

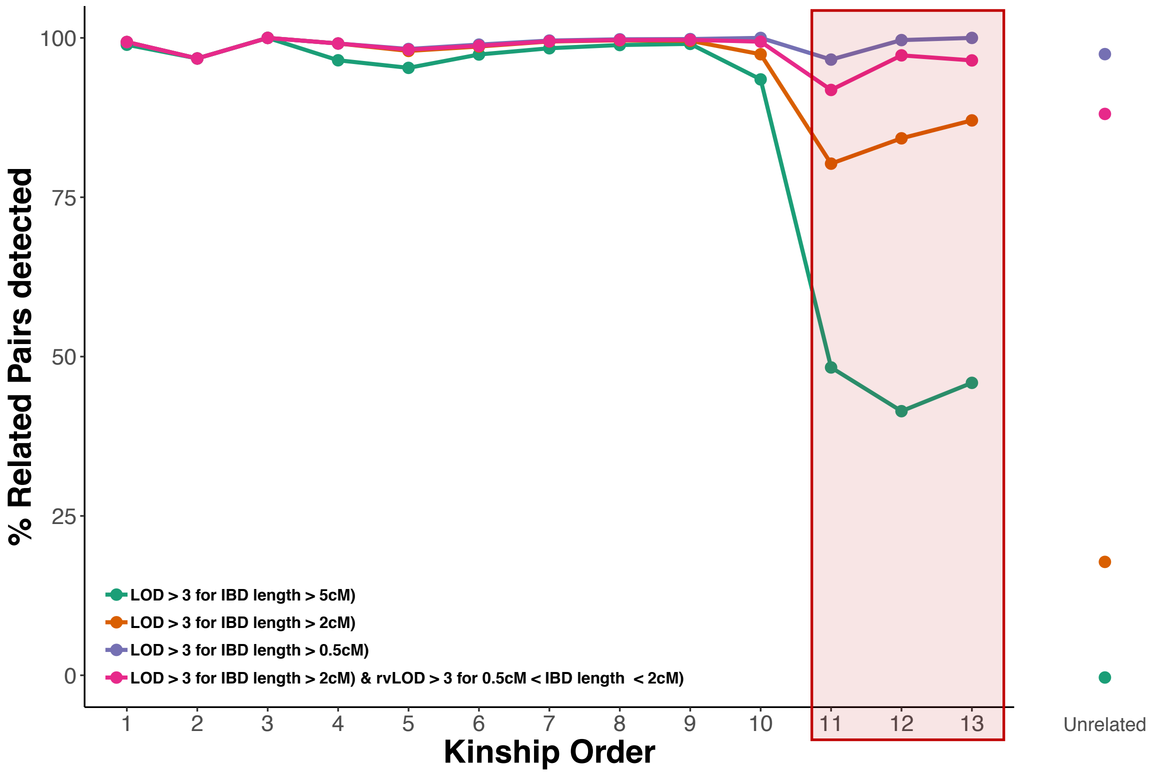


**Figure S4: Performance of rvIBD metric for relatedness estimation in the OOA cohort.**

Based on different sets of thresholds for the original log-odds (LOD) score, the enriched log-odds (rvLOD) score, and the length of IBD segment, the observed IBD proportions were compared to the expected IBD proportions. For each degree of relationship in the genotyped OOA pedigree ranging from 1st – 13th order relatedness, we computed the proportion of individual pairs with observed IBD proportions greater than the threshold set for the expected IBD proportion (see dotted line in Figure 5.5). Each line represents a different set of thresholds. The red panel highlights the higher-order (distant) relatedness where the rvIBD metric improves relatedness detection by 10% and more compared to standard IBD summary statistics.

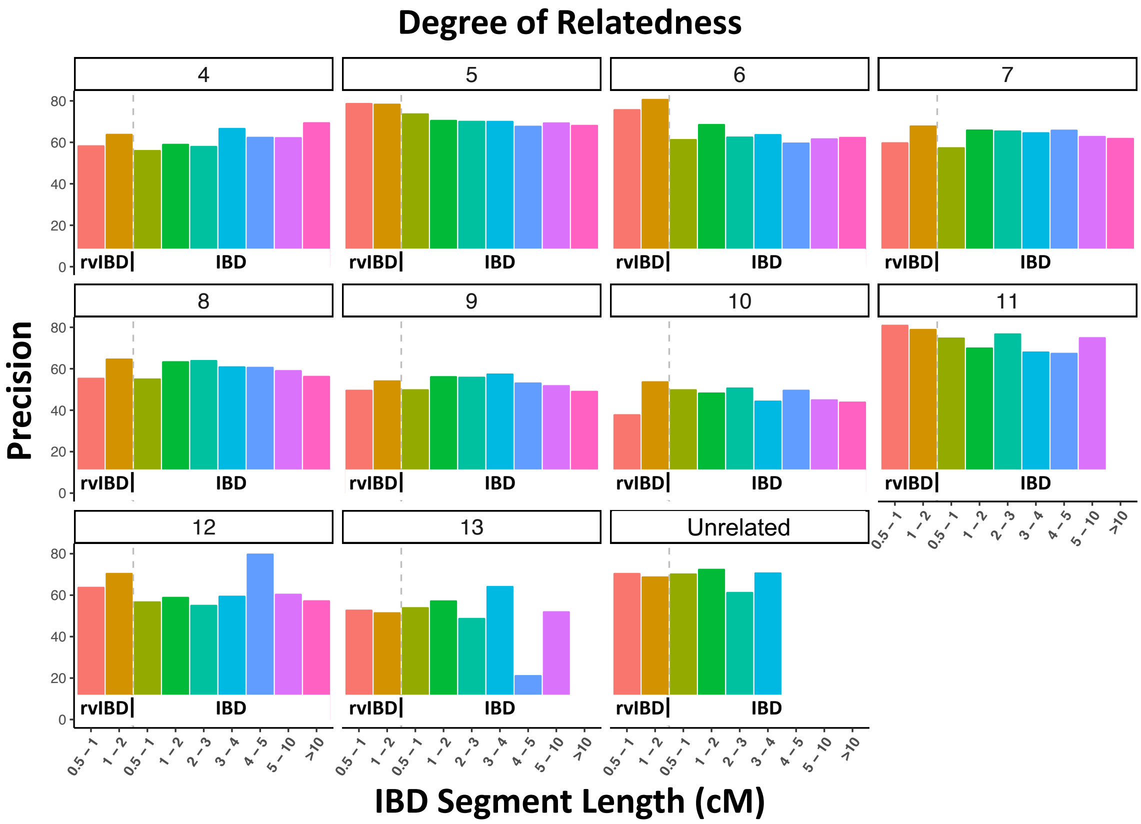


**Figure S5: Precision estimation of short IBD segments (<2cM) using the proposed rvIBD metric.**

For each degree of relationships ranging from 4th – 13th order relatedness, we computed the precision of detection of short IBD segments (<2cM) based on the rvIBD and made comparisons to the precision of detection of IBD segments segregated into bins of varying lengths based on the original IBD summary statistics.
